# A simple and reliable PCR-based method to differentiate between XX and XY sex genotypes in *Cannabis sativa*

**DOI:** 10.1101/2025.05.02.651914

**Authors:** Ainhoa Riera-Begue, Matteo Toscani, Afsheen Malik, Caroline Dowling, Susanne Schilling, Rainer Melzer

**Affiliations:** UCD School of Biology and Environmental Sciences and UCD Earth Institute, University College Dublin, Ireland

**Keywords:** Cannabis sativa, CAPS, dioecious, hemp, sex chromosome, sex determination, sex marker

## Abstract

*Cannabis sativa* is mainly a dioecious plant, which means that female and male flowers develop on separate individuals, which is controlled by an XY sex determination system: females have two X chromosomes and male plants carry an X and a Y chromosome.

*C. sativa* is a crop that has a wide variety of applications, some of which depend on the sex of the plant. Females are, for example, used for the cannabinoid production, as cannabinoids are produced in the female inflorescence. However, while adult *C. sativa* individuals present high sexual dimorphism, it is not possible to phenotypically distinguish male and female plants before flowering.

Here we present the identification of a highly conserved sex marker, *CsPDS5*, and the development of a robust, reliable and affordable PCR-based method to determine the sex genotype. In contrast to other sex genotyping methods, our approach relies on a gene polymorphic between the X and Y chromosomes and therefore requires only a single PCR with one pair of primers. The method was tested in 14 cultivars and 6 crossings, with different tissues and developmental stages, on a total of 600 samples, with 100% accuracy. Our assay allows early sex identification and hemp selection, which is useful for both research and industrial purposes.

Finally, the pipeline presented here to identify genes polymorphic between the X and Y chromosomes can serve to discover new sex markers, not only in *C. sativa*, but other dioecious plants and other organisms with 2 sexes.

## INTRODUCTION

Most flowering plants are bisexual, meaning flowers have both male and female reproductive organs. However, about 6 % of the angiosperms develop male and female flowers on separate individuals, a phenomenon called dioecy (Renner, 2014). *Cannabis sativa* is mainly a dioecious plant, although there are some monoecious cultivars with female and male flowers on the same individual (Moliterni *et al*., 2004). *C. sativa* is diploid with a pair of sex chromosomes; hence, sex is primarily determined by an XY chromosome system, female and monoecious plants are XX, and males are XY (Moliterni et al., 2004).

*C. sativa* is a multipurpose crop with a wide spectrum of uses, ranging from textiles and building materials that derive from the stalks, biofuel and oil from the seeds, to medicinal and recreational purposes thanks to the cannabinoids present in the female flower’s trichomes (Schilling *et al*., 2020). Depending on the intended use, female plants are preferred, e.g. for pharmacological applications, or male plants, e.g. for industrial applications due to their high-quality fibre content (Salentijn *et al*., 2019). Adult *C. sativa* individuals show a high degree of dimorphism (Petit *et al*., 2020), which makes it easy to differentiate between male and female plants phenotypically. However, at early stages of development before the onset of flowering, this sexual dimorphism is not present (Shi et al., 2024). Thus, for research and industrial purposes, there is a need for reliable methods to sex genotype the plants before flowering.

In recent years, different methods of sex genotyping have been developed. In most cases, MADC (male-associated DNA Cannabis) sequences have been used as markers with different strategies (Techen *et al*., 2010; Toth *et al*., 2020; Torres *et al*., 2022), some of them with great accuracy. However, it must be taken into account that MADC sequences are only associated with male individuals, presumably because they are located on the Y chromosome. Used as PCR markers, they therefore only result in amplification of male (Y chromosomal) DNA, and an autosomal control is always needed for the female or monoecious genotypes lacking a Y chromosome. Furthermore, MADC6 and some other markers are located in retrotransposons, so they may not be present in all cultivars, potentially reducing the marker’s reliability (Toth *et al*., 2020). Prentout *et al*. 2025 tried to solve this problem by identifying Y-linked genes. However, also in this assay, autosomal controls were required.

Hence, there is still a need to identify a universal genetic marker for sexing *C. sativa* seedlings. Here we present an affordable, reliable and robust PCR-based CAPS (Cleaved Amplified Polymorphic Sequence) method to differentiate between XX and XY genotypes in *C. sativa* across cultivars, as well as a pipeline that can be used to identify markers in other dioecious species.

## RESULTS

At the time of this study, no male *C. sativa* genome assembly was publicly available. The only available genome from a male individual, “JL_father” (GCA_013030025.1), was assembled at the scaffold level, making it impossible to easily identify Y chromosomal sequences.

To obtain sequences which likely originated from the Y chromosome, we previously described a computational pipeline (Shi *et al*. 2025). The pipeline uses RNA-seq data and hence does not rely on genome assemblies containing a Y chromosome. Briefly, we used RNA-seq samples from male and female *C. sativa* plants from the same developmental stage and used the following criteria to identify putative Y chromosomal transcripts. Transcripts were required to:

1. not map to the female CBDRx (GCA_900626175.2) genome.
2. have a sequence signature (kmer of 16 bp) shared by all male samples but different from all female samples.
3. be male-biased in their expression.

The pipeline resulted in the initial identification of 379 transcripts putatively originating from the Y chromosome (Shi et al., 2025).

To build on the Shi et al., 2025 method and further reduce the inclusion of false positives, a strict TPM expression level filter (average TPM > 10 in male samples, and TPM = 0 in all female samples) was applied. This allowed us to identify 25 transcripts that originated from the Y chromosome with relatively high certainty (Supplementary Data S1). From those transcripts, the genomic sequence was retrieved through a BLAST search of the ‘JL_father’ assembly. Even though this genome is not assembled at the chromosome level, blasting the mRNA sequences of the 25 putatively Y chromosomal transcripts allowed us to identify scaffolds that are most likely part of the Y chromosome. The genomic sequences of the 25 genes were then aligned to putatively corresponding sections on the X chromosome identified through a BLAST search of the CBDRx genome assembly (GCA_900626175.2). The genes with significant sequence divergence between X and Y that were deemed the most suitable for a PCR assay to distinguish XX and XY genotypes were: FE.chrY.t5, FE.chrY.t9, FE.chrY.t122, FE.chrY.t25, FE.chrY.t42 and FE.chrY.t47 (Supplementary Data 2, transcript naming after Shi *et al*. 2025). Of those, we focussed on FE.chrY.t9.

FE.chrY.t9 is homologous to the *PRECOCIOUS DISSOCIATION OF SISTERS 5* (*PDS5*) gene from *Arabidopsis* and was therefore renamed *CsPDS5*. Although both the X and Y chromosome versions of *CsPDS5* are diverged, there is a highly conserved region, which allowed the design of a common pair of primers that amplified a 419 bp region from the X as well as from the Y chromosome. Importantly, the Y version contains an *Eco*RI enzyme restriction site inside the amplicon, which is not present in the X allele. Thus, a CAPS assay in which the PCR products are subjected to a digestion with *Eco*RI results in a 157 bp and a 262 bp band for Y chromosome amplicons, while X chromosome amplicons remain at their original 419 bp length (Figure 1). Hence, visualisation via an agarose gel results in only one band at 419 bp for female and monoecious (XX) samples, while the male (XY) samples display three bands, one corresponding to the amplicon derived from the X chromosome at 419 bp and two smaller bands at 157 bp and 262 bp, corresponding to the digested amplicon derived from the Y chromosome (Figure 1). We term this assay *CsPDS5-CAPS* hereafter.

**Figure 1.**
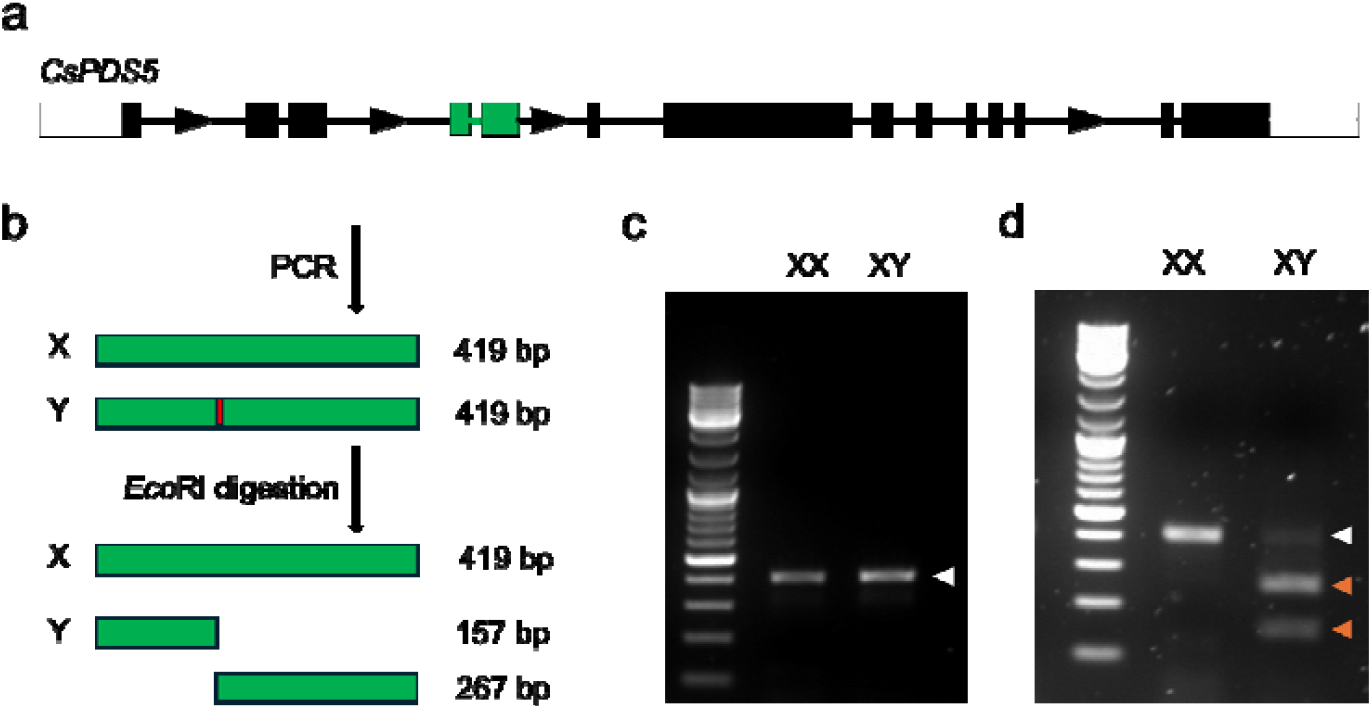
Overview over *CsPDS5-CAPS* for the *C. sativa* sex genotyping. A schematic representation of *CsPDS5* with exons and introns depicted as boxes and lines, respectively (a). The direction of transcription is indicated by arrows. The principle of the CAPS PCR-based genotyping showing amplicons in green and the *EcoRI* digestion site in red (b). Agarose gel electrophoresis of a female (XX) and male (XY) sample pre (c) and post (d) digestion. White and orange arrowheads indicate undigested (419 bp) and digested (157 and 262 bp) amplicons, respectively.

The assay was initially tested using the dioecious cultivar ‘FINOLA’. Leaf samples collected at the flowering stage, when sexual dimorphism was fully apparent, were used to easily verify that the sex as determined by *CsPDS5-CAPS* agreed with the phenotypic sex. Subsequently, ‘FINOLA’ samples at different developmental stages (from germination to flowering) and from different tissues (cotyledon and leaf) were tested, and the sex as determined by *CsPDS5-CAPS* was compared to the phenotypic sex. In total, 312 ‘FINOLA’ samples were tested, and the *CsPDS5-CAPS* genotyped sex matched the phenotypic sex in all cases.

Subsequently, we assessed the performance of *CsPDS5-CAPS* for 14 different cultivars, six dioecious (‘FINOLA’, ‘Kompolti’, ‘CSR-1’, ‘CFX-2’, ‘Estica’ and ‘Enectarol’) and eight monoecious (‘Felina 32’, ‘Bialobrzeski’, ‘Santhica 27’, ‘Earlina 8 FC’, ‘Fedora 17’, ‘Ferimon’, ‘Futura 75’ and ‘Henola’), testing two to 22 plants per cultivar. For all dioecious cultivars, our genotyping distinguished male and female individuals (Figure 2a). Monoecious cultivars are typically reported to carry two X chromosomes (Faux *et al*., 2014) and consequently appeared genotypically female in our assay (Figure 2b).

**Figure 2.**
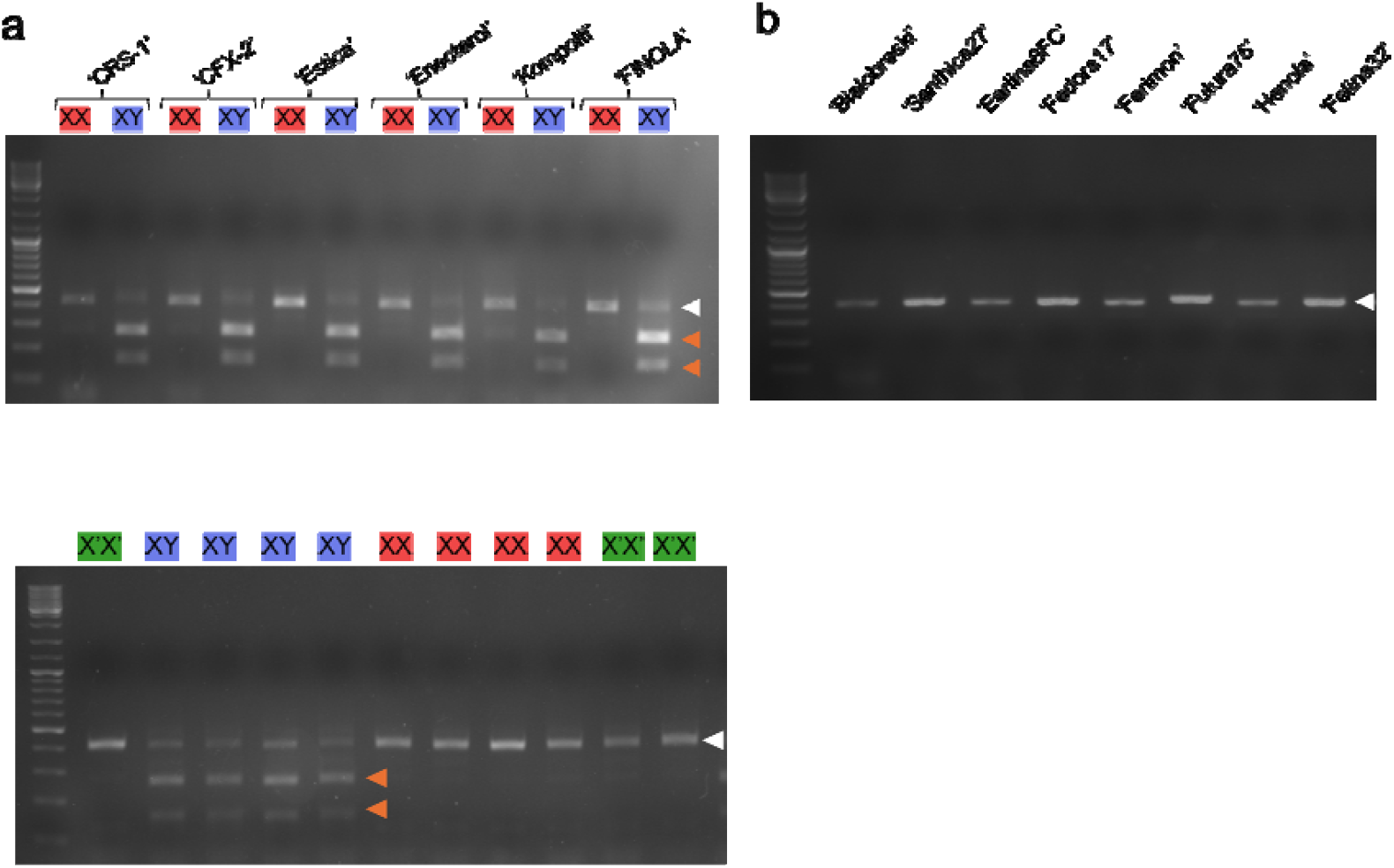
*CsPDS5-CAPS* genotyping results for different cultivars of *C. sativa*. Genotyping for female (XX) and male (XY) samples from six different dioecious cultivars (a), eight different monoecious cultivars (b) and 11 representative F2 individuals of a Finola x Felina 32 cross (c). Cultivar names and phenotypic sex are depicted above the gel image (X’X’ = Monoecious (green), XX = Female (red) and XY = male (blue)). Gel images depict PCR fragments after EcoRI digestion. Size marker Gene ruler ladder mix (Fermentas). White and orange arrowheads indicate undigested (419 bp) and digested (157 and 262 bp) amplicons, respectively.

To test the suitability of *CsPDS5-CAPS* for breeding projects, we further tested the assay on progeny plants resulting from crosses between different hemp cultivars. For this purpose, 137 F2 individuals from a previously established ‘FINOLA’ x ‘Felina 32’ cross (Dowling *et al*., 2024) were tested. As ‘FINOLA’ is dioecious and ‘Felina 32’ is monoecious, the F2 population comprises male, female and monoecious individuals (Dowling *et al*, 2024). In all 137 cases tested, our genotyping results were in agreement with the phenotypic sex (Figure 2c). We next tested F1 individuals from six additional crosses (‘FINOLA’ x ‘Earlina 8 FC’, ‘FINOLA’ x ‘Ferimon’, ‘FINOLA’ x ‘Santhica 27’, ‘FINOLA’ x ‘Fedora 17’ and ‘Felina 32’ x ‘Estica’). Also in this case, the *CsPDS5-CAPS* results were in perfect agreement with the phenotypic sex.

In total, over 500 different samples were used to test our genotyping assay. In all of the cases, the genetic and phenotypic sex matched, showing that the *CsPDS5-CAPS* is a highly reliable assay and produces robust results independently of the cultivar, crossing, developmental stage or tissue.

We next analysed whether the *CsPDS5* polymorphism utilised in our assay is also conserved in other hemp and marijuana cultivars that were not tested by PCR here but for which the genome sequences are available. We aligned the X and Y chromosomal versions of *CsPDS5* regions from different cultivars: the marijuana cultivars ‘Ace High 3-2’ and ‘SourDiesel’, the high CBD hemp cultivars ‘Pink Pepper’ and ‘GoldenRedwood’ and a US hemp landrace variety named ‘BooneCounty’ (Todd *et al*., 2024 *Preprint*). The alignment shows that the *CsPDS5* primer binding sites are conserved in all sequences, while the *Eco*RI restriction site in *CsPDS5* is present on all analysed Y-chromosomal but absent in X chromosomal sequences (Figure 3). This demonstrates that the selected region is extremely conserved across the different cultivars and that the assay introduced here is potentially useful for a wide variety of *C. sativa* cultivars and landraces.

**Figure 3.**
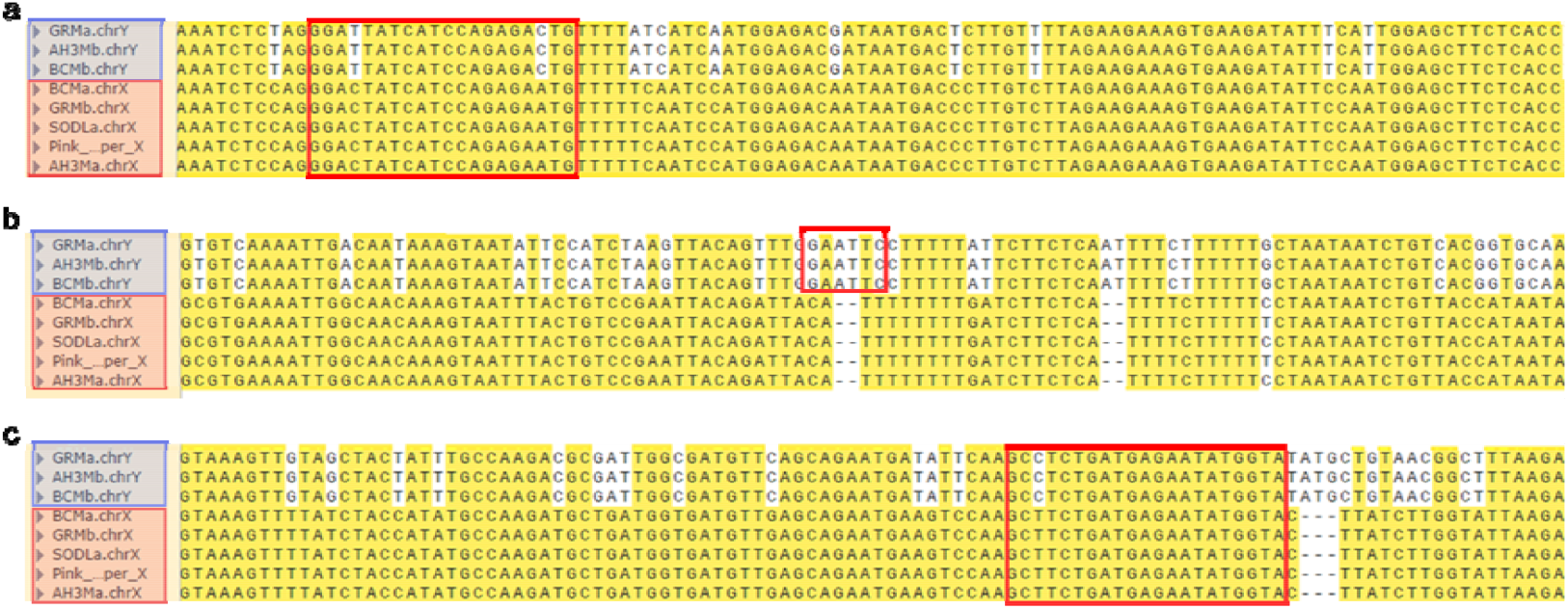
*CsPDS5* sequence from different cultivars of *C. sativa*. Alignment of three non-consecutives regions of *CsPDS5*. The red frame denotes the forward primer binding site (a), the *Eco*RI restriction site (b) and the reverse primer binding site (c). ChrX and chrY sequences from C. sativa cultivars ‘GoldenRedwood’ (GRM), ‘Ace High 3-2’ (AH3M), ‘BooneCounty’ (BCMB) and ‘SourDiesel’ (SODL) cultivars, along with with the X chromosome version of ‘Pink Pepper’ as reference genome cultivar. Bases conserved in the majority of sequences have a yellow background.

## DISCUSSION

In this study, we described the identification of a highly conserved sex marker gene, *CsPDS5*, in *C. sativa* and the development of an affordable, robust and reliable PCR-based method that is consistent among cultivars, crosses, tissue and developmental stages.

The genotyping method consists of amplifying and digesting a region of the *CsPDS5* gene, which has an enzyme restriction site on the Y version of this gene that is absent on the X version. The assay only requires the PCR reagents, a thermocycler and the EcoRI enzyme, which makes it an extremely affordable method, as only basic instruments and reagents, present in every molecular biology laboratory, are needed.

The pipeline (Figure 4) used here to identify the sex marker genes does not require an assembled Y chromosome, typically one of the more challenging tasks in genome assembly. Instead, gene expression data from male and female individuals are used to infer Y chromosomal transcripts, and genomic data are then leveraged to identify X and Y chromosomal gene copies. This basic workflow should apply to other species (plants and animals) with sex chromosomes and may therefore serve as a blueprint to identify molecular markers for sex in a range of different species. Although we leveraged the presence of a male plant assembly to retrieve the transcripts corresponding Y chromosome genomic sequence, the method can be applied also to species missing genomes assemblies if exploiting transcript polymorphisms instead of introns polymorphisms.

**Figure 4.**
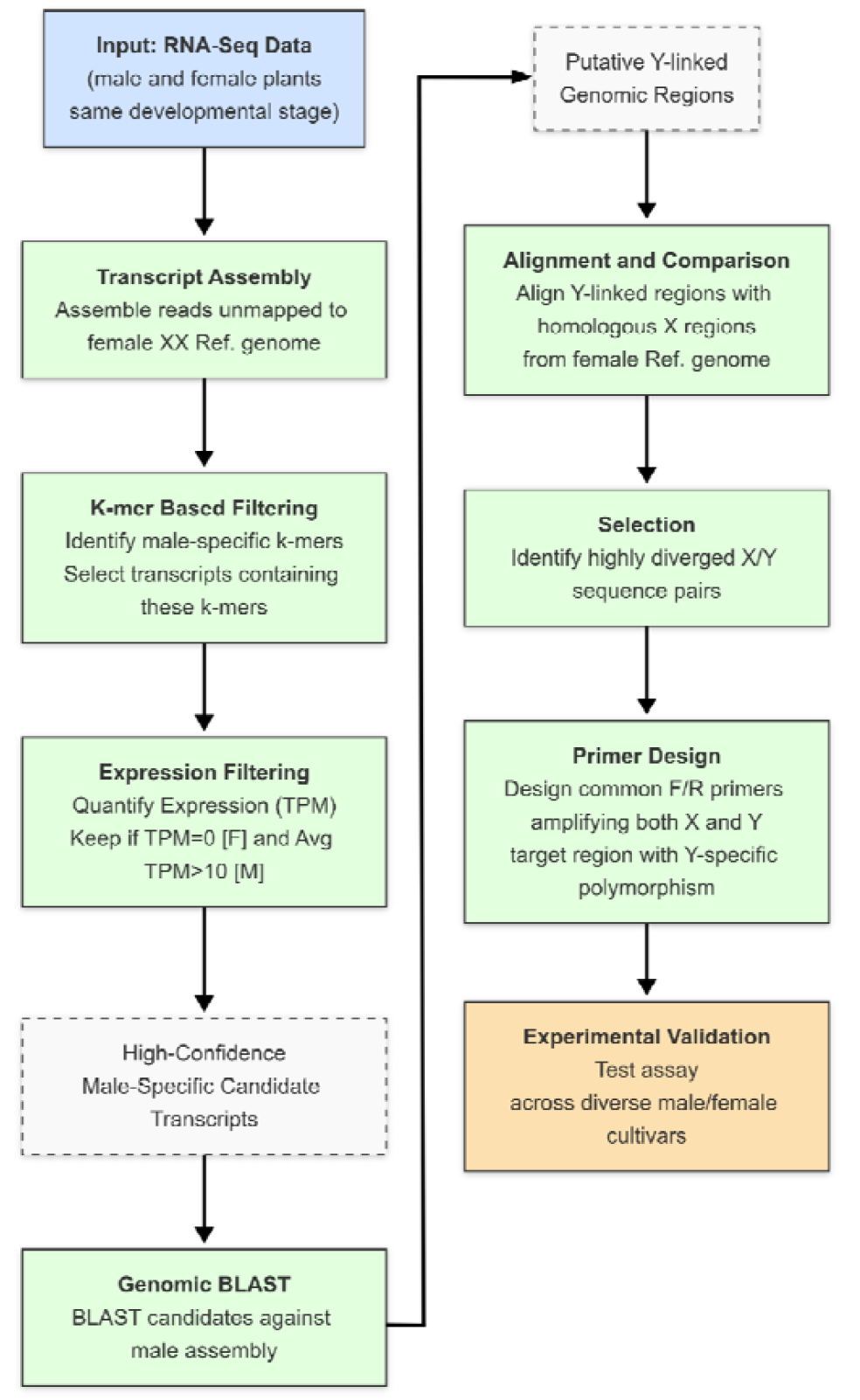
Pipeline used to identify molecular sex markers

We have tested the *CsPDS5-CAPS* assay on more than 500 samples from 14 cultivars, six crossings, different tissues and developmental stages, and in all of the cases, the genetic and phenotypic sex matched, showing an accuracy of 100 %. Analyses of publicly available genomes further demonstrated the presence of the *Eco*RI restriction site and of the and primer binding sites also in other hemp and marijuana cultivars, indicating that the *CsPDS5* polymorphism is well conserved in *C. sativa*, probably due to its involvement in an essential role in the function of the cohesin complex (Panizza, et al. 2000). Other *C. sativa* sex markers, like MADC6, are located in retrotransposons and may have experienced copy number variations and transposition (Toth *et al*., 2020). *CsPDS5* might be a functional gene and thus not be subject to the same constraints as MADC6.

Also, since *CsPDS5* is present in both the X and the Y chromosomes, it simultaneously provides information about XX and XY genotypes using a single primer pair, which is different from other sex genotyping methods (Techen *et al*., 2010; Toth *et al*., 2020; Torres *et al*., 2022; Prentout, *et al*. 2025).

Our assay was developed to provide a robust sex genotyping method using basic laboratory equipment. However, it should be noted that, since the basis of the assay is a 3 bp difference between X and Y chromosomal *CsPDS5* copies (Figure 3), the marker should also be suitable for high throughput methods such as TaqMan SNP genotyping assays or PACE, which could be used in larger breeding programs. Likewise, amplicon sequencing should be equally suitable for the *CsPDS5* polymorphism.

The sex expression of *C. sativa* is known to be influenced by various factors, and the relative contribution of genetic vs. environmental factors for sex expression remains somewhat unclear (reviewed in Schilling *et al*., 2020). The observation that our molecular sex marker was in complete agreement with the phenotypic sex for hundreds of samples, including a mapping population that originated from a cross of a dioecious with a monoecious cultivar, illustrates that genetic components play a very strong role in sex determination in *C. sativa*. Unquestionably, external factors like the application of silver nitrate, an ethylene inhibitor that leads to the development of male flowers on female plants, can modify the sex expression (Galloch, 1978; Truta, *et al*., 2007). However, our observations indicate that under conventional growth conditions, sex is almost solely determined by the XY sex determination system.

## MATERIAL AND METHODS

### Gene selection

The pipeline to identify Y chromosomal transcripts is outlined in (Shi *et al*., 2025). This pipeline leverages RNA-seq data from male and female individuals, comparative kmer analysis, and filtering criteria to isolate candidate Y-linked genes with higher confidence (Shi *et al*., 2025).

Briefly, the first step of the method is to assemble the transcripts from the reads that fail to map to the reference female XX chromosome genome, as at least part of those are likely to originate from the diverged genes on the Y chromosome. Then, to further refine the resulting transcripts from false positives, we employed a k-mer-based method selecting only the transcripts containing 16-mers present in every male sample, but absent in every female sample. Finally, the transcripts are increasingly refined by quantifying their expression level and selecting the ones showing a statistically significant male bias, with filtering parameters of log2 fold change (log2FC) ≥1 and false discovery rate (FDR) P-value ≤0.05.

To further reduce the possibility of false positives, here we selected as candidate genes for the genotyping assay only transcripts presenting a TPM level of 0 in females and a relatively significant expression in males (mean TPM > 10).

After identifying male-specific sequences, the whole genomic sequence was inferred by BLASTing the mRNA sequence against the JL Lion father genome (Genome ID: GCA_013030025.1). As a last step, the whole genomic region, comprising introns, from the reference genome chromosome X and JL Lion chromosome Y, was aligned, and the genes showing significant divergence in sequence between their chromosome X and chromosome Y versions were selected as the most suitable candidates.

### DNA isolation

The DNA was isolated from the corresponding sample using the DNeasy® Plant Mini Kit (Qiagen GmbH, Germany) according to the manufacturer’s instructions. DNA quality was checked by electrophoresis in a 1% agarose gel at 120 mV for 20 minutes, and DNA quantity was determined using a Nanodrop spectrophotometer (ND-1000).

### PCR primer design and conditions

Both versions of the *CsPDS5* gene, from the X and the Y chromosome, were aligned using Clustal Omega and this alignment was used to manually design the primer pair (Table 1), the specificity of which was checked against the NCBI database using BLASTn tool. The primers amplify a fragment of 419 bp for both versions of the gene. The primers were purchased from IDT. The IDT primer stocks were then used to create diluted stocks with concentrations of 50 µM.

**Table 1.**
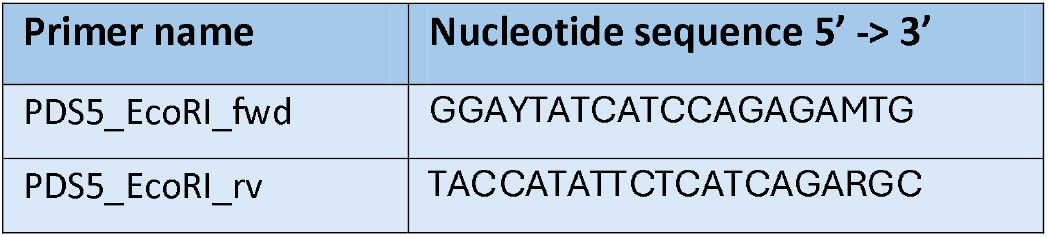
Name and nucleotide sequences of the genotyping primers both forward and reverse.

PCRs were conducted in a final volume of 25.5 μL with: 17.75 μL nuclease-free water, 5 μL 5X Phusion HF Buffer (Thermo Fisher Scientific) which provides 1.5 mM MgCl2 in the final 1X concentration), 0.5 μL of dNTPs (10 mM each) (VWR Chemicals), 0.5 μL PDS5_EcoRI_fwd (50 μM), 0.5 μL PDS5_EcoRI_rv (50 μM), 0.25 μL of Phusion DNA Polymerase (2 U/μL) (Thermo Fisher Scientific) and 1 μL of DNA template (≥ 2ng/μL).

The Biometra T3000 Thermocycler was used for the PCR. After an initial denaturation at 98 °C for 30 s, 40 cycles at 98 °C for 10 s, at 56.5 °C annealing temperature for 30 s, and an extension at 72 °C for 30 s were performed before a final extension at 72 °C for 5 min.

### *Eco*RI Digestion

After the PCR, the products were subjected to a digestion with the restriction endonuclease *Eco*RI which recognises the sequence 5’-GAATTC-3’. The digestion reaction consisted of 10X 1.3 μL of FastDigest Green Buffer (ThermoFisher Scientific), 1 μL FastDigest EcoRI (12.5 ThermoFisher Scientific), 12.5 μL of the unpurified PCR product and 4.5 μL of nuclease-free water. The mix was incubated at 37°C for 15 minutes.

### Gel electrophoresis

The digestion products were loaded on a 2 % agarose gel containing SYBR safe DNA gel stain (invitrogen) and run for 30 minutes at 100 mV. The gel was then visualised under UV.

## ACKNOWLEDGMENTS

We thank Darren Dougharty for help with the PCR genotyping. This publication has emanated from research conducted with the financial support of Taighde Éireann - Research Ireland under grant number in this publication was funded by the Irish Research Council under grant number IRCLA/2022/3294.

## DATA AVAILABILITY

All relevant data can be found within the manuscript and its supporting material.

## SUPPORTING INFORMATION

Data S1: 25 putatively Y chromosomal transcripts.

Data S2: Fasta alignment files of 5 candidate genes potentially suitable as sex markers in *C. sativa*.

